# Early exposure to PFAS disrupts neuro-muscular development in zebrafish embryos

**DOI:** 10.64898/2026.01.19.700343

**Authors:** Zainab Afzal, Brian N. Papas, Vandana Veershetty, Evan E Pittman, Charles Hatcher, Jian-Liang Li, Warren Casey, Deepak Kumar

## Abstract

Development is a tightly regulated process that establishes body axes and orchestrates the spatial organization of tissues and organs. Although developmental programs contain inherent redundancies, they remain highly sensitive to environmental cues. Among environmental contaminants, per- and polyfluoroalkyl substances (PFAS), chemicals that resist degradation and bioaccumulate in the body, are of particular concern. These “forever chemicals” are widespread in our household products, including non-stick and waterproof materials, and drinking water remains a major source of exposure. PFAS accumulate in specific tissues and have been associated with developmental delays, childhood leukemia, and other adverse health outcomes, yet the cellular and molecular mechanisms by which they disrupt early development remain largely unknown. To address this, we employ zebrafish embryos as a New Approach Methodology (NAM) to investigate how perfluorooctanoic acid (PFOA), a prevalent environmental PFAS, alters early embryogenesis. Embryos were exposed to physiologically relevant low and high doses of PFOA and analyzed at 24 hours post-fertilization (hpf), a key stage of organogenesis. We also included a parental exposure group, in which adults were treated with PFOA and their offspring were collected to assess whether the effects of exposure were transmitted to the next generation. Developmental processes are inherently plastic, and we wanted to understand the extent to which PFOA impacts normal cellular processes as well as the redundancy in the system (different developmental signaling pathways) which ensures that an embryo develops properly. Towards this, we performed single-nucleus RNA sequencing at 24 hpf, and it revealed that neuronal and muscle tissue clusters are particularly sensitive to PFOA exposure. These molecular perturbations correspond with anxiety-like behavioral phenotypes we observed in the exposed larvae, linking early developmental disruptions to organism-level outcomes. Overall, our findings provide mechanistic insight into the way in which PFAS exposure alters development, disrupting gene expression patterns and chromatin organization in developing tissues, revealing how early molecular perturbations can give rise to long-term behavioral consequences.

**Graphical Abstract:** 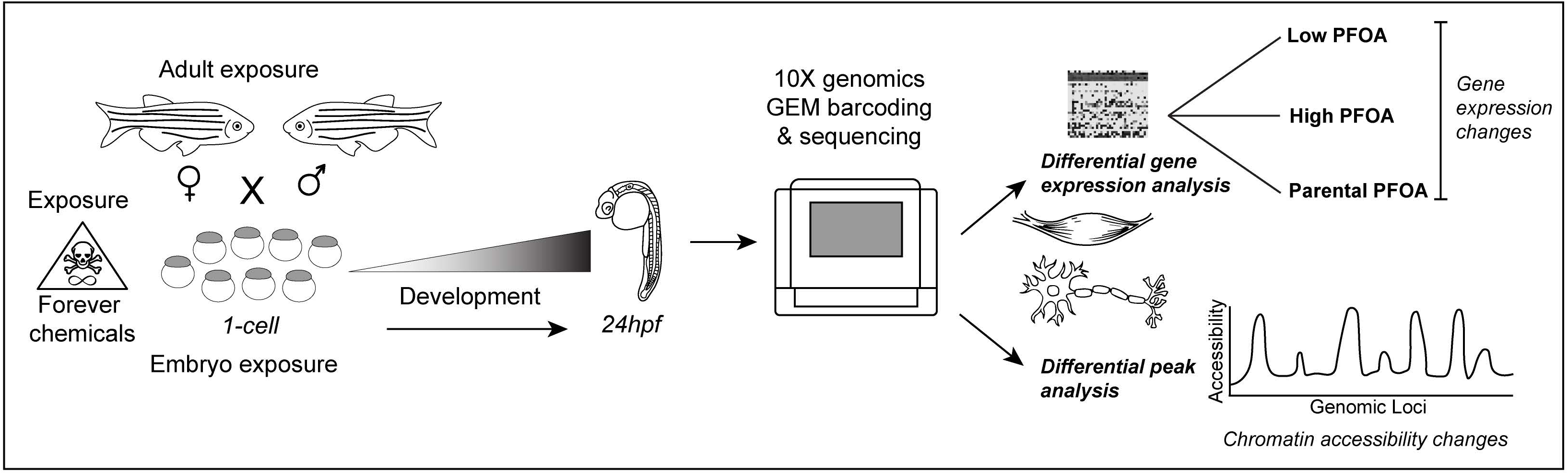

## Introduction

Development of an embryo involves fundamental gene regulatory programs that establish body axes, direct the spatial patterning, and drive cellular differentiation, organization, and architecture of organs to ensure the proper functioning of an organism. Although these developmental programs are robust and often buffered against perturbations, they remain sensitive to external environmental cues during critical windows of development. Disruptions during these periods can alter the execution of developmental gene networks, leading to aberrant tissue patterning, delayed maturation, or long-term physiological and behavioral consequences. Natural instances of response to environmental cues impacting morphology of an organism or developmental outcomes have also been observed. For example, variations in temperature can impact which butterfly morph will grow from the larvae (Koch & Bückmann, 1987; Nijhout, 1991). Sex determination in reptiles also depends on temperature instead of chromosomes, such that eggs incubated below 30°C produce females while eggs incubated above 34°C produce males (Ferguson & Joanen, 1982). Oher examples include variations in nutrient availability, oxygen tension, or mechanical forces which can alter the activity or distribution of morphogens and thereby impact tissue patterning, growth, and organogenesis (Fathollahipour et al., 2018; Hannezo & Heisenberg, 2019; LeGoff & Lecuit, 2015). These examples illustrate how environmental inputs can intersect with core developmental processes that govern tissue identity and function.

In contrast to natural environmental variation, exposure to environmental toxicants can negatively impact developmental processes, often resulting in acute or chronic disease. For instance, exposure to lead, arsenic, mercury have been associated with adverse impacts on neurocognitive development (Kaur et al., 2025), while alcohol and corticosteroids have been associated with muscle toxicity and altered myogenesis (Alleyne & Dopico, 2021; Pereira & Freire de Carvalho, 2011). These findings underscore the vulnerability of neuronal and muscle tissues, systems that emerge early in development and are essential for coordinated motor and behavioral function.

Beyond naturally occurring toxicants, exposure to anthropogenic chemicals such as per- and polyfluoroalkyl substances (PFAS), commonly referred to as “forever chemicals,” have emerged as a growing public health concern. Owing to their resistance to degradation, PFAS have become ubiquitous environmental contaminants, widely detected in water, air, and soil, and are increasingly recognized as sources of physiological disruption (Fenton et al., 2021/03; Hall et al., 2023; Plunkett, Application July 1, 1939,; Xu et al., 2020/12/01). PFAS exposure has been linked to multiple adult disease etiologies including adult kidney and liver disease (DeWitt et al., 2019; Fenton et al., 2021/03; Qi et al., 2023; Rosen et al., 2022; Salihovic et al., 2019; Schmidt, 2022/05; Sunderland et al., 2019). Additionally, epidemiological studies have also associated PFAS exposure with developmental delays, including language deficits and reduced motor scores in toddlers, as well as decreased lean muscle mass in teenagers with higher exposure levels (DeLuca et al., 2023; Eick et al., 2023; Janis et al., 2021; Jones et al., 2023; Kingsley et al., 2019; Luo et al., 2022). Despite these associations, the molecular mechanisms by which PFAS specifically disrupt embryonic neuronal and muscle development remain largely unexplored.

A central challenge in understanding PFAS developmental toxicity lies in the inherent redundancy and compensatory capacity of developmental programs. While gross morphology even after toxic exposure may appear preserved, environmental toxicants may exert highly cell-type-specific effects that get masked in bulk analyses. Subtle perturbations in transcriptional regulation, chromatin accessibility, or spatial gene expression within exposure vulnerable cell populations may nevertheless lead to persistent functional deficits later in life. Dissecting these effects therefore requires approaches capable of resolving developmental responses at single-cell resolution and within their spatial context.

The zebrafish (*Danio rerio*) is a powerful vertebrate model for elucidating the mechanisms of PFAS toxicity during early development. Zebrafish embryos develop externally, are optically transparent, and produce large numbers of synchronized progeny, enabling direct visualization of developmental processes and high-throughput experimental designs. The rapid emergence of functional neuronal and muscle systems within 24 hours post-fertilization further makes zebrafish well suited for interrogating early neuromuscular development. Moreover, 70% of the zebrafish genome is conserved with humans (Howe, 2013), facilitating direct comparisons of genetic and regulatory pathways between zebrafish and humans. Collectively, these features establish zebrafish embryos as a New Approach Methodology (NAM) (Ball et al., 2025), and they are already widely used in chemical hazard evaluation through the Zebrafish Embryotoxicity Test (ZET) framework (Achenbach et al., 2020). With regard to assessing PFAS toxicity, because PFAS readily dissolve in water, environmentally relevant exposure paradigms can be implemented simply by treating embryos in their aquatic environment, enabling high-throughput assessment of developmental toxicity.

In this study, we integrate developmental phenotyping with single-nucleus transcriptomic and chromatin accessibility analyses to determine how early-life exposure to perfluorooctanoic acid (PFOA), a commonly found type of PFAS, disrupts embryonic development at cellular and molecular resolution. We first assess morphological and physiological endpoints during early development, followed by single-nucleus profiling to uncover tissue-specific transcriptional responses. Our analyses reveal pronounced gene expression changes, particularly within neuronal and muscle cell populations. Moreover, PFOA exposure alters chromatin accessibility at and around genomic regions associated with these dysregulated genes, implicating epigenomic disruption as a mechanistic link between early environmental exposure and subsequent neuromuscular and behavioral outcomes. Consistent with these molecular alterations, exposed larvae exhibit altered behavioral phenotypes. Together, our findings provide a comprehensive mechanistic framework linking PFAS exposure to disrupted developmental gene regulation, tissue patterning, and organism-level consequences.

## Results

### Early-life PFOA exposure effects on morphological and physiological development

To assess whether PFOA, an eight-carbon chain PFAS still ubiquitous in our environment, affects zebrafish embryonic development, we treated embryos at the 1-cell stage with low and high concentrations of PFOA (Fig. 1A). All 1-cell treatments were performed within 5–10 minutes of collection to ensure exposure began within 15 minutes, before embryos reached the 2-cell stage.

**Figure 1:**
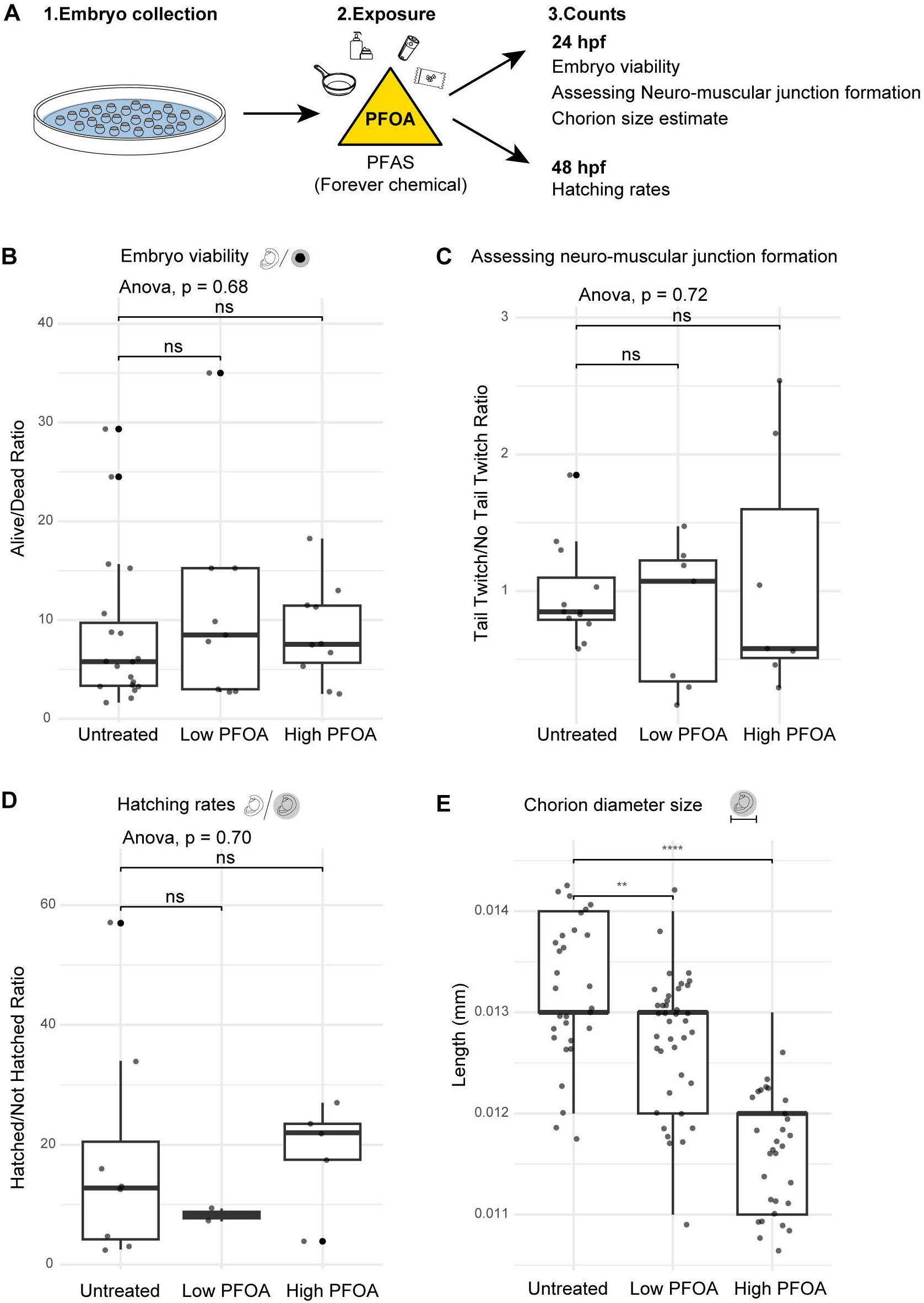
PFOA exposure impacts developmental traits in zebrafish embryos. A - Zebrafish embryonic exposure schematic to depict timing of exposure and recordings of viability and behavior. B - Embryo viability assessment with ratio of alive over dead for each exposure condition at 24hpf. C - Assessment of neuromuscular junction formation by manually counting tail twitch response after a poke, done at different times and by 3 different people to account for any natural variations. D - Hatching ratio plotted by counting how many embryos hatched out of the chorion at 48hpf vs how many didn’t hatch for each exposure and untreated condition. E - Group images were taken for treated and untreated embryos along with a microscopic scale. Measurements of chorion diameters were done for each exposure and untreated.

Embryos were allowed to develop, and at 24 hours post-fertilization (hpf), we quantified viability by counting dead or developmentally arrested embryos relative to normally developed embryos. Apart from an occasional developmental arrest, PFOA exposure did not significantly increase abnormal development. Interestingly, while not statistically significant, the average ratio of alive to dead embryos was slightly higher in treated groups compared to controls across different experimental days, suggesting a trend toward increased viability (Fig. 1B).

Given prior evidence that PFOA affects neuronal development, we next performed a rapid behavioral assay to determine the lowest exposure at which neuromuscular function was altered. Embryos were treated with concentrations ranging from 50 ng/L to 10,000 ng/L and assessed at 24 hpf for tail twitch in response to a tactile stimulus, indicative of functional neuromuscular junctions (Supp. Fig. 1). The lowest concentration showing a detectable difference was 700 ng/L (or 0.00169 μM), which we selected as our low exposure dose. The high dose (25 μM or 1,040,000 ng/L) was chosen based on environmental and human exposure data, representing still a low environmental concentration in some areas but a high serum level in humans. Using zebrafish embryos, which provide a rapid and efficient system to assess multiple exposure conditions by simply adding different concentrations of PFOA to the water, we assessed proper neuromuscular junction function by recording tail twitches as our quick readout in response to tactile stimulus. While zebrafish embryos naturally exhibit spontaneous tail twitching around 24 hpf, we specifically recorded twitches elicited by a mechanical poke for our behavioral assay. Although the tail twitch responses were not statistically significant, a trend emerged with low PFOA exposure tended to increase tail twitch frequency, whereas higher exposure reduced responsiveness (Fig. 1C). This pattern suggests a dose-dependent effect on neuromuscular development, with higher exposures potentially delaying maturation of neuromuscular junctions and reducing the number of embryos capable of responding to tactile stimulation.

We further assessed developmental progression via observing hatching rates between 48–72 hpf, recording whether PFOA had any effect on an embryos ability to hatch out of its chorion. While no significant differences were observed between control and treated groups, a trend was noted such that low exposure resulted in fewer hatched embryos, while high exposure had slightly more hatched embryos than controls (Fig. 1D). This trend again hints towards dose dependent changes occurring with PFOA exposure.

In addition to hatching rates being an indicator of normal development rate, it has been observed that changes in chorion sizes can be used as a measure of response to toxicity and even there have been changes in somatogenic mutants exhibiting smaller chorion sizes (Pérez-Atehortúa et al., 2023). We wanted to check if PFOA, which is trending towards impacting normal development is having an impact on chorion sizes. We observed that the diameter for the chorion was significantly decreased with exposure to PFOA, with a higher exposure amount having an even smaller chorion diameter (Fig 1E). This reduction appeared to occur within the first hour of exposure (Supp. Fig. 2), raising the possibility that PFOA affects chorion size through osmotic effects, mechanical forces, or embryo-mediated physiological responses. Early embryonic mechanical forces are known to influence development (Villeneuve et al., 2025), suggesting that PFOA-induced changes in chorion diameter could potentially have significant downstream effects on embryogenesis.

### Single-nuclei resolution of PFOA-induced effects during embryogenesis

To investigate the cellular mechanisms underlying the developmental disruptions observed with PFOA exposure, including potential developmental delays and morphological changes such as reduced chorion sizes, we examined the extent to which PFAS affects gene expression and chromatin organization at the single-nuclei level. We focused on early life exposure to capture molecular changes during key developmental stages when organogenesis has occurred or been initiated, specifically at 24 hpf.

We also assessed whether parental exposure influenced embryonic development in the absence of direct embryonic exposure (Fig. 2A). For all exposure conditions, embryos were allowed to develop until 24 hpf and then flash-frozen. Nuclei were simultaneously isolated from all samples to minimize batch effects and processed using the 10X Multiome workflow (Fig. 2B), enabling simultaneous analysis of gene expression and chromatin accessibility at single-cell resolution.

**Figure 2:**
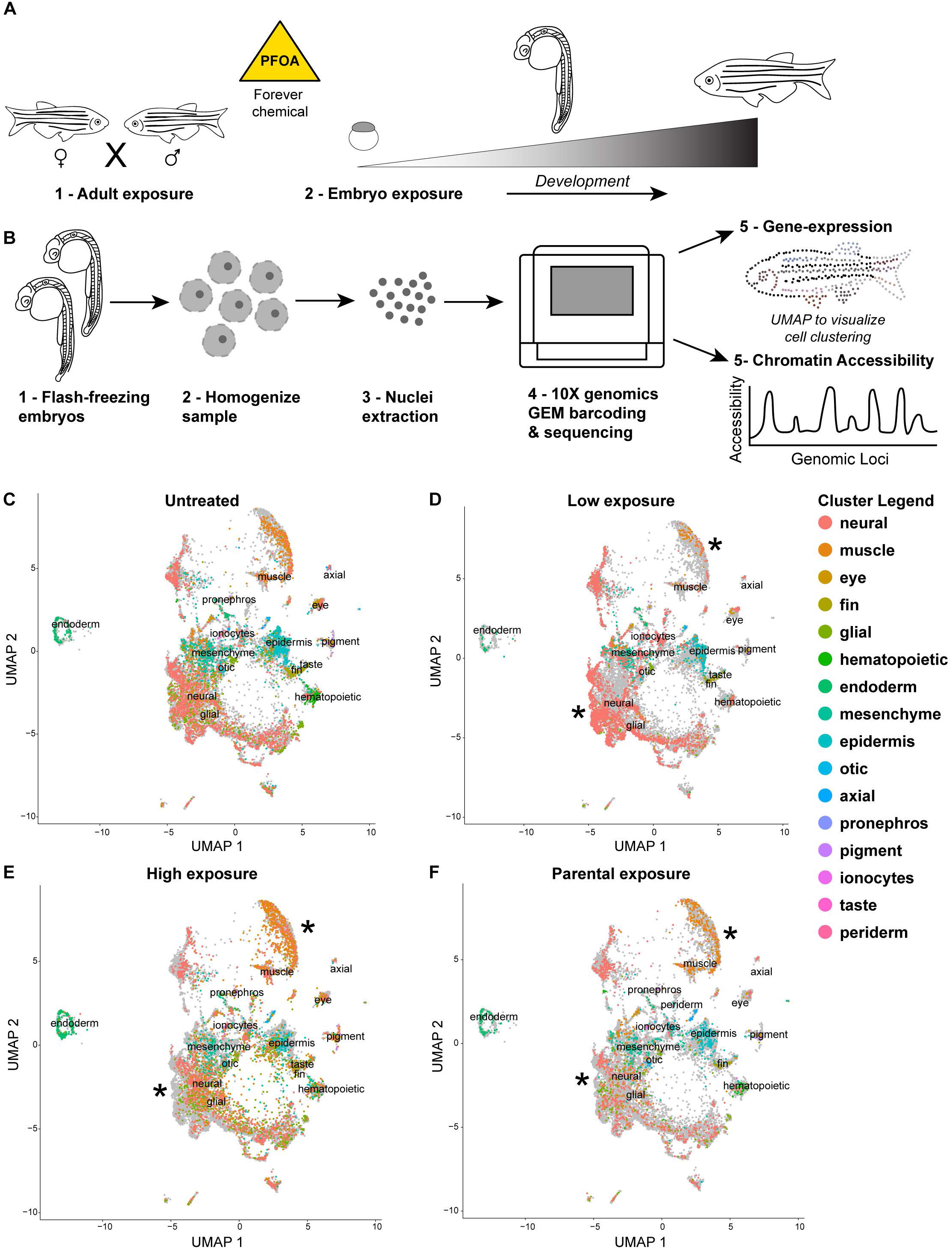
Single nuclei resolution of developmental disruption with PFOA exposure. A – Schematic illustrating the PFOA exposure paradigms for embryos and adults. B – Workflow showing how embryos that were flash-frozen at 24 hpf were processed for single-nucleus isolation and subsequently prepared for 10x Multiome profiling to obtain both gene expression and ATAC-seq datasets. C-F – Feature maps showing the cell clusters identified across untreated and PFOA-exposed samples. Each color represents a distinct tissue type, as indicated in the cluster legend. Asterisks (*) denote tissue clusters that exhibit the most pronounced changes under exposure conditions.

All samples were visualized using a combined feature or expression plot to compare treated embryos to untreated controls (Fig. 2C–F). To better understand what cell types were changing in expression patterns when exposed to PFAS, tissue cluster identities were assigned using the Daniocell single-cell zebrafish database (Sur et al., 2023). With this approach we identified 16 distinct tissue types across our samples. This allowed for comparison of gene expression patterns across tissue clusters in response to PFAS exposure.

To compare across the tissue clusters, we also normalized for cell counts to see how expression of specific genes was changing in the different tissues. We first looked at the clusters that have the greatest transcriptional changes across exposure treatments compared to untreated and saw that in most cases select neuronal and muscle clusters are the most effected tissue cluster types with exposure to PFOA (supp fig3). The epidermal cluster also exhibited alterations, although whether PFOA of any PFAS can traverse the skin and directly influence epidermal cells remains to be determined. To more efficiently compare gene expression changes across clusters, we compared heatmaps for statistically significant differential genes for all the different tissue clusters (supp fig4-5) and compared the most differential expressed genes within the tissue clusters. The expression patterns of genes across the muscle and neuronal clusters are significantly changing in the exposure conditions, and it appeared that there is a difference in response based on the dosage of PFOA.

In the low exposure condition, the number of cells represented for the muscle cluster appear to be reduced but it is not significantly different (supp fig6), while in the high exposure condition, the number of muscle cells appears to increase. Maybe this represents that the dosage of PFOA during development impacts the number of cells that are differentiating into muscle. Thus, it appears that the higher exposure is causing more cells to differentiate into muscle cells, maybe as a compensation mechanism for not reaching the right muscular activity level. Regardless of changes in the tissue densities, when normalizing for cell counts, and then comparing for gene expression changes across the tissue clusters, it still appears that the muscle tissue cluster is significantly impacted with PFOA exposure.

In the neuronal cluster, the trend in cell densities was reversed compared to muscle cells. In the low exposure condition, there is higher density of neuronal cells in the tissue cluster while there is a reduction in density with the higher exposure. Does this potentially indicate that more neurons are being differentiated in the low exposure and could serve as an early indicator of anxiety related behavior issues? While these observations could suggest dose-dependent effects on neuronal differentiation, behavioral outcomes are probably also dependent on additional factors such as neuronal wiring and synapse formation. Importantly, after normalizing for cell counts, several gene expression patterns within the neuronal cluster were significantly altered in response to PFOA exposure.

Parental PFOA exposure also influenced embryonic gene expression, even in the absence of direct exposure. In this exposure condition, adult fish were housed in water containing low dose of PFOA, but mating tanks were PFOA-free, and collected embryos were allowed to develop in standard embryo media. While changes across tissue clusters were less pronounced than in directly exposed embryos, neuronal and muscle clusters remained significantly affected, suggesting that parental exposure impacted gametes or altered epigenetic marks which were inherited by the offspring. Neuronal cell counts were comparable to controls, whereas muscle cell counts were significantly increased, consistent with a potential compensatory mechanism or epigenetically driven shift in differentiation. Further studies, including direct tracking of PFAS bioaccumulation in specific tissues, are needed to correlate exposure levels with cellular responses. Currently, no effective methods exist to visualize PFAS distribution within embryonic tissue clusters in vivo.

### PFOA exposure reshapes muscle and neuronal gene expression patterns

To determine trends in gene expression across exposure conditions, we compared heatmaps of the top differentially expressed tissue clusters, focusing on muscle and neuronal populations (Fig. 3A). Although fold-change values were generally modest across all exposure groups, this likely reflects our use of a lower, environmentally relevant dose, which enhances sensitivity for detecting subtle developmental disruptions. Several genes were differentially expressed across multiple conditions, while some exhibited changes unique to a single exposure. In the muscle cluster, most top differentially expressed genes were downregulated relative to controls. In the neuronal cluster, both up- and downregulated genes were observed. Notably, some genes showed dose-dependent changes; for example, *Foxp4* was downregulated at low PFOA exposure but upregulated at high exposure. Parental exposure induced fewer differential expression changes, and no genes were differentially expressed across all conditions in neuronal tissue.

**Figure 3:**
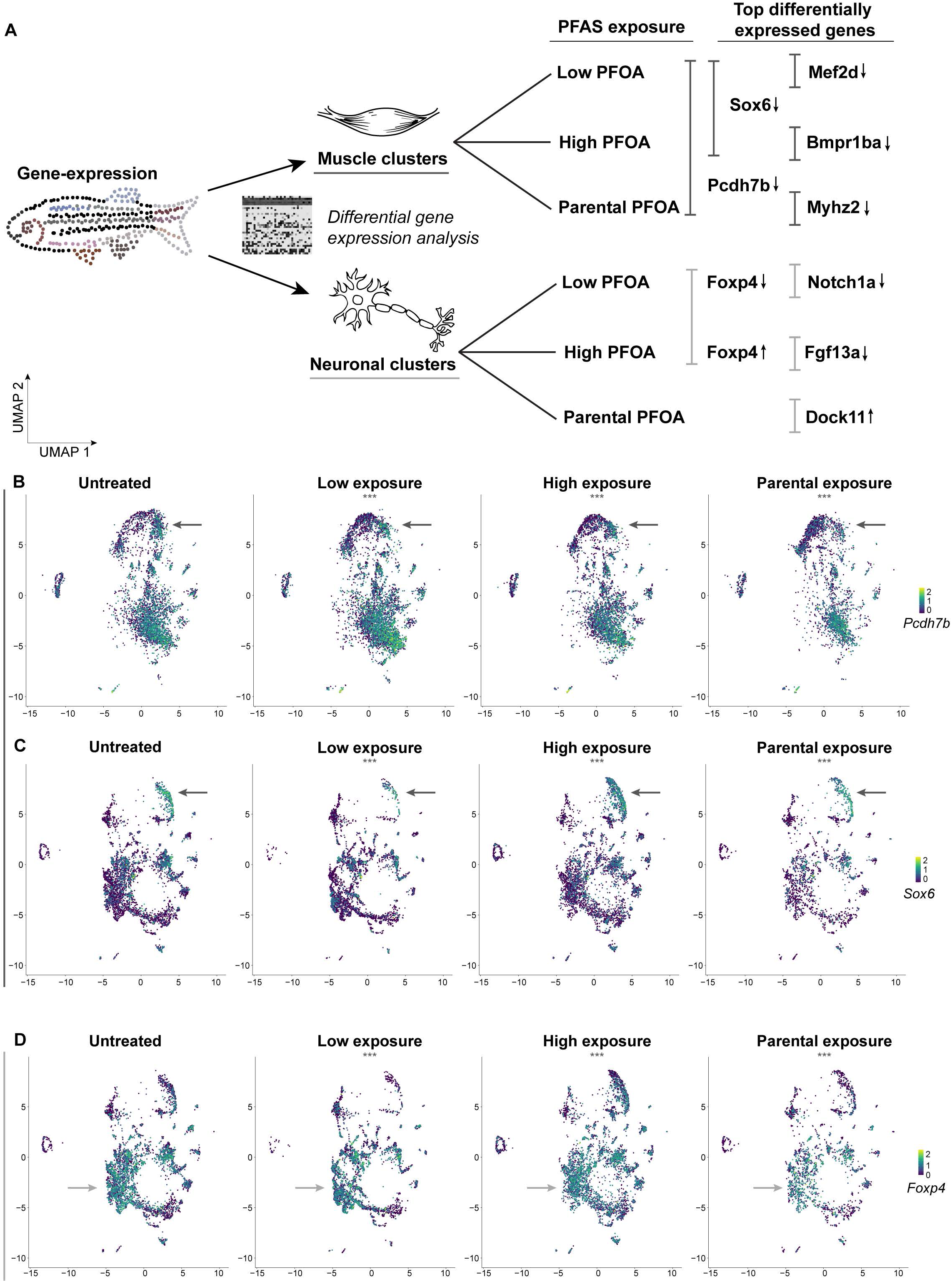
PFOA exposure effect on the neuronal and muscle clusters during development. A – Schematic summarizing differential gene expression changes identified in the 10x Multiome dataset, highlighting the most affected neuronal and muscle clusters. Selected top differentially expressed genes are shown with arrows indicating whether they are upregulated or downregulated. B-C – UMAP feature plots showing expression of *pcdh7b* and *sox6*, two of the most differentially expressed genes in muscle tissue, displayed across all clusters and exposure conditions. Arrows point towards the muscle cluster. D – UMAP feature plots showing expression of *foxp4*, a gene differentially expressed in the neuronal cluster, displayed across all clusters and exposure conditions. Arrows point towards the neuronal cluster.

While we found instances of several genes that were changing across conditions, one gene consistently affected across both embryonic and parental exposures was *pcdh7b*, which was downregulated in the muscle cluster (Fig. 3B). *Pcdh7b* is one of five zebrafish protocadherins, highly conserved with human orthologs, with distinct expression patterns in the developing central nervous system, including the forebrain, midbrain, neural tube, and myotome (part of a somite that will develop into a muscle and is usually innervated by a single spinal nerve) during development(Blevins et al., 2011; Thisse et al., 2004). Functionally, PCDH7 has been implicated in cell migration, where its polarized expression is thought to promote movement by facilitating phosphorylated myosin complexes and ERM proteins, which crosslink actin filaments to plasma membranes, thereby establishing the necessary mechanical properties for cell migration (Qureshi et al., 2022). PCDH7 has also been implicated in sarcopenia, the progressive loss of muscle mass, strength, and function (Liu et al., 2023). Specifically, in our dataset, downregulation of *pcdh7b* was most pronounced in slow muscle fibers (annotations from the Daniocell database), fibers which are critical for long-term endurance relying on aerobic muscle activity (Barresi et al., 2001; Kuznetsov et al., 2025). This observation raises the possibility that PFAS exposure may impact muscular function, particularly in slow muscle fibers involved in endurance activity. Given that aerobic exercise has been shown to reduce anxiety levels (Broman-Fulks et al., 2004), these findings suggest a potential link between PFAS-induced changes in *pcdh7b* expression and alterations in anxiety-related behaviors.

We also observed downregulation of *Sox6* in the same slow muscle fiber cluster. *Sox6* is a transcription factor (TF) known for its role in repressing slow fiber differentiation. Knockout studies have indicated that reduced *Sox6* expression increases slow muscle fibers and increases mitochondrial activity, enhancing endurance (Quiat et al., 2011). Analysis of the *pcdh7b* genomic locus revealed SOX6 binding sites (Supp. Fig. 7), suggesting a potential disruption in a connected cis-regulatory mechanism. In humans, SOX6 mutations have been linked to developmental delays including intellectual disability, attention-deficit/hyperactivity disorder (ADHD), autism, mild facial dysmorphism, and craniosynostosis (Tolchin et al., 2020). All together, our data indicate that PFAS exposure may alter transcriptional regulatory programs in slow muscle fibers, potentially affecting functional muscle development. These early transcriptional changes were also detected in precursor somatic mesoderm tissue, with significant downregulation of both *pcdh7b* and *Sox6* in low and high exposure conditions (Supp. Table 1). This highlights that PFAS-induced disruptions emerge early and persist as cells differentiate into slow muscle fibers.

Neuronal clusters were similarly affected. Dose-dependent changes in *Foxp4*, a gene implicated in neuronal development (Del Viso et al., 2023), were observed across neural subtypes. Low exposure decreased *Foxp4* expression in general neurons, neuronal progenitors of the midbrain and hindbrain, and the diencephalon, whereas high exposure increased expression in hindbrain, diencephalon, ectoderm, and telencephalon subtypes. These region-specific and dose-dependent effects underscore the importance of exposure level during early neural development, as affected regions contribute to forebrain structures such as the thalamus, hypothalamus, posterior pituitary, and pineal gland, which are critical for homeostasis and organ function.

Collectively, these results demonstrate that PFOA exposure alters transcriptional programs in both muscle and neuronal clusters in a dose-dependent manner, with effects observed at early developmental stages and in differentiated cell populations. These findings provide insight into molecular mechanisms by which PFAS exposure may contribute to neuromuscular and neurodevelopmental phenotypes, highlighting the importance of considering total dose of exposure during early development.

### Spatial dynamics of target genes in the developing zebrafish

Having observed significant gene expression changes in muscle, we next sought to spatially characterize these patterns in developing muscle using fluorescent *in situ* hybridization via the hybridization chain reaction (HCR) technique (Fig. 4A). We focused on the 24 hpf zebrafish tail, a region where neuromuscular junctions are forming. Probes were designed against *pcdh7b*, *Sox6*, and *Foxp4*, and all three genes were visualized simultaneously in the same embryo.

**Figure 4:**
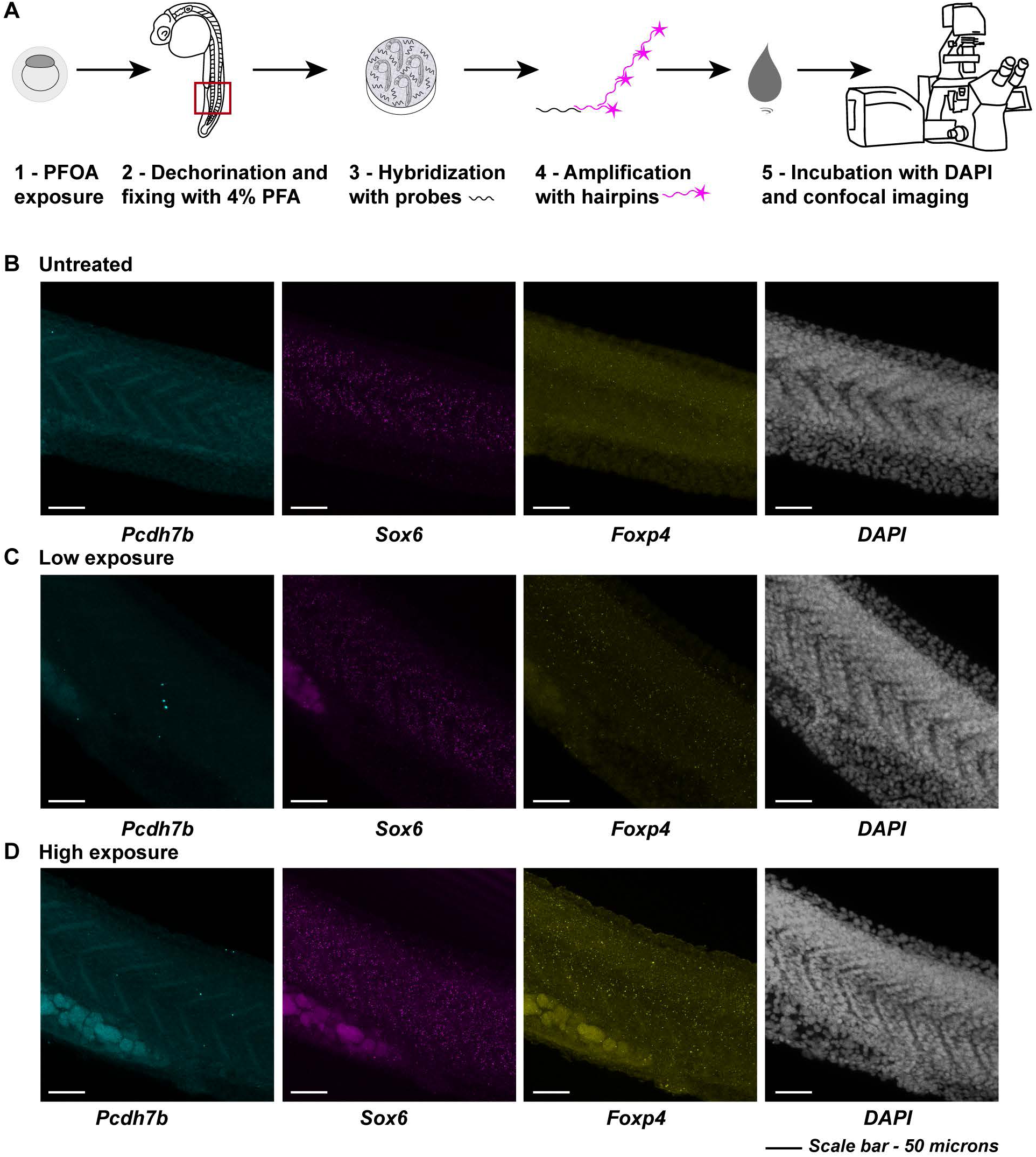
Spatial localization of differentially expressed muscle and neuronal cluster genes during development in response to PFOA. A - Schematic illustrating the Hybridization Chain Reaction (HCR) in situ workflow used to fluorescently visualize spatial gene expression patterns in developing embryos. B – Expression of *pcdh7b* (cyan), *sox6* (magenta), and *foxp4* (yellow), along with DAPI, in the developing zebrafish tail (region indicated by a red square in panel 4A-2) in untreated embryos. C - Expression of *pcdh7b* (cyan), *sox6* (magenta), and *foxp4* (yellow), along with DAPI, in the same tail region in embryos exposed to a low dose (700 ng/L) of PFOA. D - Expression of *pcdh7b* (cyan), *sox6* (magenta), and *foxp4* (yellow), along with DAPI, in the same tail region in embryos exposed to a high dose (25 µM) of PFOA.

In untreated embryos, *pcdh7b* expression was localized to the marginal zone of the spinal cord, visible as V-shaped structures between somites along the tail. Expression decreased under both PFOA exposure conditions, with low exposure showing a greater reduction (Fig. 4B–D, left panels), consistent with observations from our single-nuclei dataset. To quantify *pcdh7b* transcript abundance, we used the Imaris machine-learning algorithm. Because we observed few bright nascent transcriptional puncta and predominantly single processed transcripts, we quantified total summed intensity across the entire *pcdh7b*-expressing region to capture overall expression changes. This analysis revealed a similar pattern of reduced *pcdh7b* expression with PFOA exposure (Fig. 5A). These results are further supported by quantification of *pcdh7b* expression normalized to the number of DAPI-positive nuclei (Fig. 5A’).

**Figure 5:**
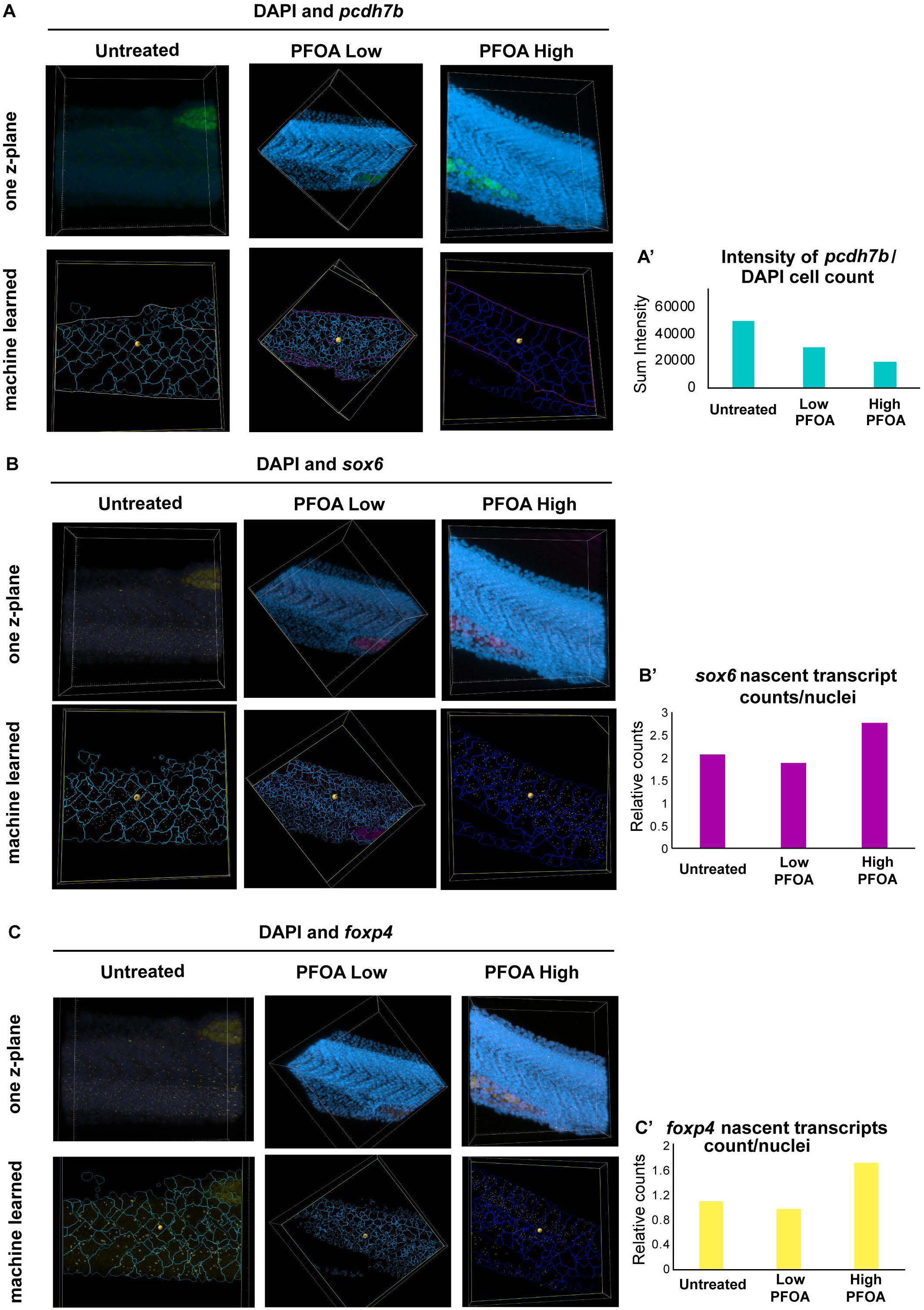
Quantifying HCR signal for each channel and comparing against DAPI using Machine learning in Imaris. A – Top panels show one z-plane to illustrate DAPI and *pcdh7b* expression, which primarily appears as single transcripts. Bottom panels show the machine-learning output in Imaris, where cell number was estimated using DAPI as the training channel, and *pcdh7b* sum intensity was calculated within regions of expression. **A′** displays quantification of *pcdh7b* sum intensity normalized to cell count across exposure conditions compared to untreated controls. B – Top panels show one z-plane of DAPI and nascent *sox6* transcripts. Bottom panels show the machine-learning output, where cell number was estimated using DAPI and the number of bright *sox6* puncta (nascent transcripts) was quantified. **B′** displays quantification of nascent *sox6* transcripts normalized to cell count across exposure conditions compared to untreated controls. C – Top panels show one z-plane of DAPI and nascent *foxp4* transcripts. Bottom panels show the machine-learning output, where cell number was estimated using DAPI and the number of bright *foxp4* puncta (nascent transcripts) was quantified. **C′** displays quantification of nascent *foxp4* transcripts normalized to cell count across exposure conditions compared to untreated controls.

*Sox6* nascent, newly synthesized transcripts were clearly visible in the somites of wild-type embryos, and this expression was significantly reduced under the low-dose exposure condition (Fig. 4B–D, middle panels). Because nascent transcripts were abundant across all conditions, we again used the Imaris machine-learning algorithm to detect these bright punctate transcriptional spots (Fig. 5B). Quantification showed that most cells expressed two nascent transcripts, indicating that both alleles are actively transcribed under normal conditions. This level of transcriptional activity decreased with low-dose PFOA exposure, consistent with the reduced expression observed in our single-nucleus dataset (Fig. 5B’). Intriguingly, we observed a marked increase in the number of nascent transcripts in the high-dose exposure condition, opposite to the trend in the single-nucleus dataset. One possibility is that cells in the high-dose group may be dividing more frequently and co-transcribing during the cell cycle. Alternatively, the apparent increase in nascent transcripts could reflect compensatory transcription in response to impaired processing or degradation of processed transcripts following PFOA exposure. Further experiments will be required to distinguish between these possibilities.

For *Foxp4*, wildtype embryos displayed abundant processed transcripts with a few nascent transcripts interspersed within and around the somites. Low exposure reduced both nascent and processed transcript levels, whereas high exposure increased expression, mirroring dose-dependent trends observed in the single-nucleus dataset for neuronal expression (Fig. 4B–D, right panels). Because *Foxp4* was also affected in the diencephalon, we examined forebrain expression and observed a similar pattern, such that low exposure decreased expression while high exposure increased it (Supp. Fig. 8). In tissue sections, *foxp4* expression appeared predominantly as bright punctate nascent transcriptional spots along the tail (Fig. 5C). Quantification using the Imaris machine-learning algorithm showed that under normal conditions, cells typically contained one nascent transcript, suggesting one active allele with a cell (Fig. 5 C’). With low-dose PFOA exposure, the number of nascent transcripts decreased, indicating fewer actively transcribing cells, a trend consistent with our single-nucleus dataset. Intriguingly, high-dose PFOA exposure resulted in an increased number of nascent transcripts per cell, again aligning with our single-nucleus dataset. When normalized to cell count, many cells appeared to express two nascent transcripts, suggesting active transcription from both alleles. This raises two possibilities, one that high PFOA exposure may slow transcriptional shutoff, increasing the likelihood of capturing two active sites per cell, or two that it may trigger a compensatory upregulation of *foxp4* in response to other developmental disruptions caused by exposure. Further experiments will be required to distinguish between these possible reasons. Overall, these results validate our single-nucleus findings and demonstrate spatial, dose-dependent alterations in gene expression across developing muscle and neuronal tissues following PFOA exposure. With *sox6* and *foxp4*, by visualizing nascent transcripts at sites of active transcription, we observe clear disruptions in transcriptional dynamics, suggesting that PFOA may interfere with the transcriptional machinery within these cells of the developing zebrafish tail.

### PFOA exposure perturbs chromatin accessibility neuronal and muscle clusters during development

Given the disruption of target gene expression with PFOA exposure, we next investigated potential mechanisms underlying these changes. PFAS compounds have been reported to bind DNA (Qin et al., 2024/01/01), raising the question of whether they could bind and alter chromatin architecture *in vivo*. To address this, we analyzed single-nuclei ATAC sequencing peaks to assess chromatin accessibility across tissue clusters in the different exposure conditions (Fig. 6A). All treatment conditions exhibited different changes in chromatin accessibility, with the strongest effects observed under low and high PFOA exposure, while parental exposure had minimal impact. Compared to untreated embryos, many regions exhibited increased chromatin accessibility, particularly at the upstream and intronic regions which could act as enhancer and cis-regulatory elements controlling transcription. Increased accessibility can either facilitate or repress TF binding, but given that many target genes were downregulated, these changes may reflect recruitment of repressive factors to these open chromatin regions in the treated conditions.

**Figure 6:**
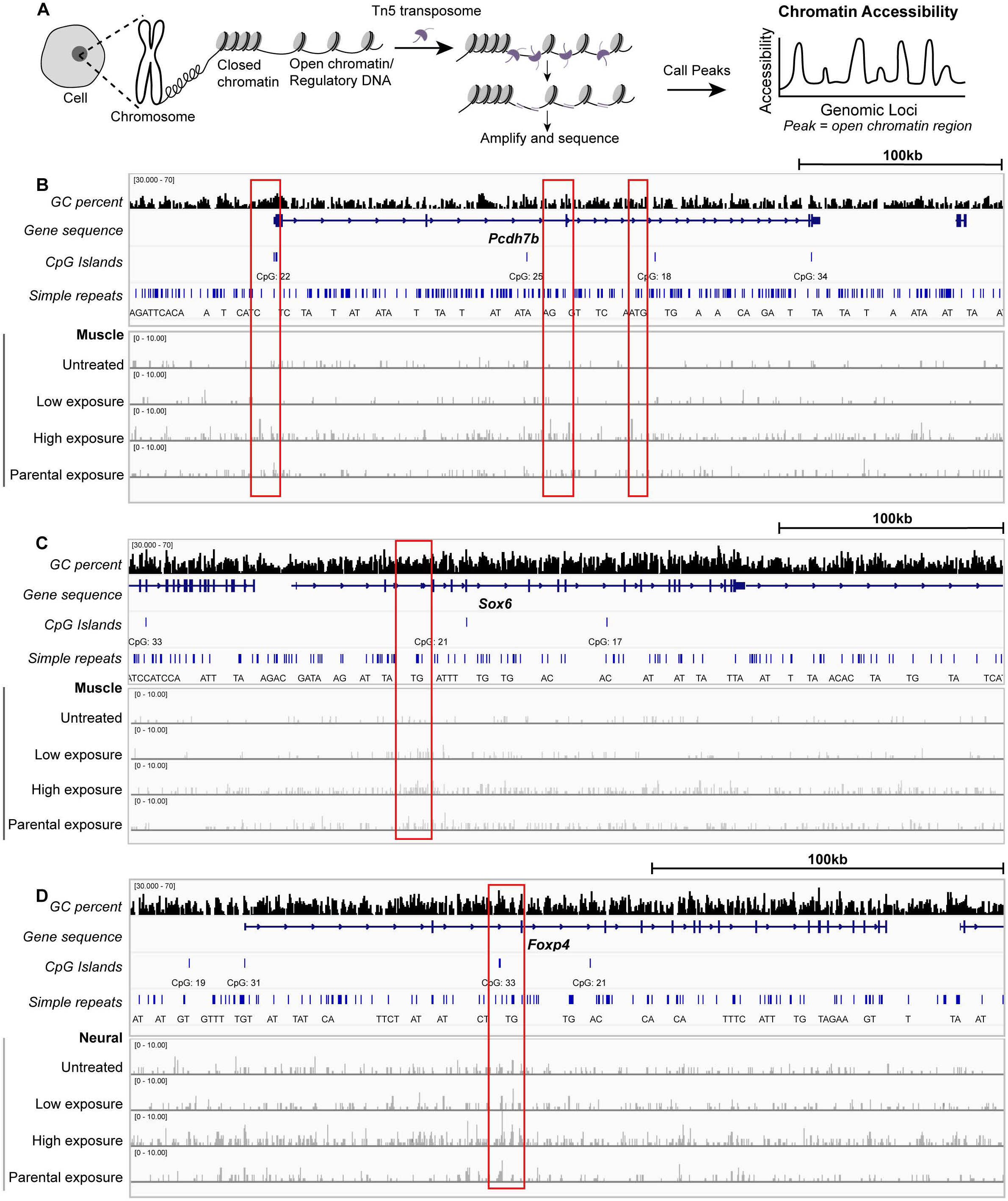
Impact of PFOA exposure on chromatin accessibility in neuronal and muscle clusters during development. A – Schematic illustrating the ATAC-seq workflow used to identify open chromatin regions (peaks) across the different exposure conditions. B – Genome browser tracks for the *pcdh7b* locus, showing gene structure, GC content, CpG islands, and simple repeats. Bottom panels display ATAC-seq profiles for the muscle cluster across exposure conditions compared to untreated embryos. C – Genome browser tracks for the *sox6* locus, showing gene structure, GC content, CpG islands, and simple repeats. Bottom panels display ATAC-seq profiles for the muscle cluster across exposure conditions compared to untreated embryos. D – Genome browser tracks for the *foxp4* locus, showing gene structure, GC content, CpG islands, and simple repeats. Bottom panels display ATAC-seq profiles for the neuronal cluster across exposure conditions compared to untreated embryos.

To further investigate potential regulatory mechanisms, we examined specific loci within target genes to identify transcription factor (TF) binding sites. For example, we evaluated whether *sox6*, a TF with predicted binding sites in the *pcdh7b* locus, to assess whether these sites overlapped with regions of altered chromatin accessibility following PFOA exposure. Although a comprehensive, tissue-specific map of all TF binding sites, and the enhancer grammar underlying their activity, is not yet available in zebrafish, this targeted approach provides initial insight into regulatory disruptions affecting transcription during development. To achieve this, we selected DNA regions that showed exposure-dependent accessibility changes (Fig. 6, red rectangles) and queried them against the catalog of inferred zebrafish TF binding preferences from CisBP (Weirauch et al., 2014). We matched all TF motifs in the database to the altered regions and identified enrichment for several TF families. The top most binding motifs present in the altered DNA regions included zinc-finger proteins (e.g., jazf1b, znf366, znf362b), forkhead factors (e.g., foxd3, foxd5), and homeobox-containing TFs (e.g., hnf1bb, dbx1b). Many of these factors are known to contribute to chromatin architecture, enhancer activity, and regulation of downstream gene transcription (Supp. Figs. 9–11; Supp. Tables 2–4). Notably, several motifs correspond to TFs involved in methylation and demethylation pathways that maintain active enhancer states, suggesting that PFOA exposure may disrupt early transcriptional programs by perturbing epigenetic regulators.

Given that proper epigenetic regulation is essential for neurodevelopmental processes, these disruptions likely contribute to the behavioral phenotypes observed later in life. Consistent with this, we observe that PFOA exposure correlates with measurable behavioral alterations in developing larvae. At 5 days post-fertilization, when larvae display robust light-dark responses, we assessed movement and responses to environmental stimuli. PFOA-exposed larvae exhibited increased locomotor activity and a preference for peripheral zones, consistent with heightened anxiety-like behavior (Supp. Fig. 12). These behavioral changes likely reflect early disruptions in neuronal and muscle clusters, suggesting that altered chromatin accessibility during development may have lasting effects on neuromuscular wiring and sensory responsiveness as the exposed embryo develops. Notably, PFOA did not impact overall viability, yet these results indicate significant functional consequences for normal developmental processes, highlighting PFOA as an environmental risk factor affecting muscular and neuronal development.

## Discussion

Persistent environmental toxicants pose significant risks to embryonic development and organismal physiology. Here, we examined how early-life exposure to the ubiquitous PFAS, PFOA, disrupts zebrafish development at both cellular and molecular levels. Early developmental phenotyping revealed subtle but measurable alterations in morphology and physiology, including significant changes in chorion sizes and trends towards delayed tail twitch response, suggesting that even environmentally relevant exposures can perturb fundamental developmental processes. These findings underscore the vulnerability of embryos during critical windows of neuromuscular development.

Single-nucleus transcriptomic analyses revealed pronounced, cell-type-specific disruptions in neuronal and muscle lineages. In neuronal clusters, PFOA altered gene expression in progenitors of the midbrain, hindbrain, and diencephalon, regions essential for brain development, motor coordination, and organ regulation. Muscle tissue, particularly slow fibers and their precursors in the somatic mesoderm, also exhibited significant transcriptional perturbations. Spatial mapping of target genes using HCR fluorescent *in situ* hybridization demonstrated disruptions in nascent transcription, indicating that PFOA interferes with gene regulatory programs in precise spatial domains within tissues. Together, these data suggest that PFOA compromises both tissue-specific transcriptional identity and regulation of transcriptional activity.

Chromatin accessibility profiling further illuminated potential mechanisms underlying these transcriptional changes. Several loci corresponding to dysregulated genes showed increased accessibility, many containing transcription factor binding sites critical for chromatin organization and tissue-specific regulation. These findings suggest that PFOA may disrupt normal chromatin remodeling, either directly or via interference with chromatin-associated factors, potentially perturbing cis-regulatory networks essential for tissue-specific gene expression. Integrating transcriptional and epigenomic data supports a model in which early molecular disruptions propagate through the developmental process, affecting tissue formation and function.

Importantly, molecular and spatial disruptions observed during embryogenesis correlate with functional consequences at later developmental stages. PFOA-exposed larvae displayed heightened anxiety-like behavior, particularly in response to stress stimulus of light. These observations challenge the assumption that PFAS-induced behavioral phenotypes arise solely from neural dysfunction, highlighting the potential contribution of muscle tissue perturbations to neuromotor outcomes. Our data suggest that early disruptions in transcriptional and chromatin programs may prime specific tissues for long-term functional deficits.

Our study emphasizes the value of single-cell and spatial approaches for understanding environmental toxicant effects. Future work employing fluorescently labeled PFAS to track bioaccumulation and determine whether these chemicals remain associated with DNA or chromatin proteins, will help to clarify whether observed effects reflect transient insults or persistent molecular challenges. Combining multi-omic profiling with functional and behavioral assays will be critical to fully connect molecular perturbations to altered tissue architecture and organismal outcomes.

In conclusion, we propose a unifying framework in which PFOA exposure during early development alters chromatin accessibility and transcriptional programs within neuronal and muscle lineages, disrupts spatial gene expression and tissue patterning, and contributes to behavioral phenotypes in exposed larvae. These findings provide a mechanistic link between environmental PFAS exposure and developmental dysregulation, highlight the susceptibility of neuromuscular systems to persistent chemical toxicants, and underscore the utility of zebrafish as a high-resolution model for developmental toxicity studies.

## Acknowledgements

We gratefully acknowledge support from the RCMI Pilot Award to ZA, funded by the RCMI grant U54MD012392. We are also grateful to support from the PFAS Network grant through the NC Collaboratory. Additionally, we would like to thank Nicole Salazar Velmeshev, Qing Cheng, Derek Norford, Vivekanandan Ramalingam, Jaya Krishnan, Younshim Park, and Khyati Dalal, Marina Venero Galanternik for their guidance, support, and insightful discussions on the project.

## Materials and Methods

### Animal Husbandry and embryo culture

Adult zebrafish (Danio rerio, wild-type AB strain) were maintained in the North Carolina Central University Animal Resource Facility. All procedures were approved by the Institutional Animal Care and Use Committee (IACUC; Protocol D16-00378, approved 06-01-2022). Adults were housed on a 14:10 h light:dark cycle in a recirculating Aquaneering system (San Diego, CA, USA) using reverse osmosis (RO) water. Embryos were maintained in embryo medium (RO water buffered with Instant Ocean© sea salt) at 28.5 °C. System pH was maintained between 7.0 and 7.5.

Adult fish were fed an irradiated diet (Zeigler Bros. Inc., Catalog #AH271) twice daily and supplemented with live brine shrimp (EG Artemia, INVE Aquaculture Inc.). Embryos for experiments were obtained by setting up adult breeders at a 2:4 male-to-female ratio in breeding tanks, with sexes separated overnight. At 09:00 the following morning, dividers were removed, and embryos were collected 15 minutes later. Collected embryos were rinsed in RO water, transferred to embryo medium containing 0.1% methylene blue, and sorted into treatment groups prior to incubation at 28.5 °C.

### PFOA exposure conditions setup

Perfluorooctanoic acid (PFOA) (Sigma, Catalog #171468-5G, ≥95% purity) was dissolved in embryo medium to prepare a 1000 µg/L stock solution. Working concentrations were prepared fresh for each experiment. Embryos were exposed to either a low dose of 700 ng/L (∼1.7 µM) PFOA or a high dose of 25 µM (∼10,400,000 ng/L). Exposures began at the 1-cell stage and embryos were allowed to develop after sorting in regular or treated embryo media at 28.5 °C in 60-mm Petri dishes (Corning Catalog #FB0875712) until 24 hours post-fertilization (hpf), at which point they were dechorionated and flash-frozen.

For parental exposure, untreated and PFOA-treated adults were maintained off-system on separate racks. Adults assigned to the treatment group were exposed to 700 ng/L PFOA and housed in 2.8-L tanks. Reverse osmosis (RO) water was changed every other day for both treated and untreated groups. After two weeks of low-dose PFOA exposure, embryos were collected from the adult fish. Breeding tanks contained only clean embryo medium (no PFOA) to ensure that collected embryos were not directly exposed to PFOA during spawning.

At 24 hpf, embryos were assessed for viability under a dissecting microscope. Dead or developmentally arrested embryos were scored as nonviable, while viable embryos were defined as those reaching the appropriate morphological stage for 24 hpf. Tail-twitch responses were also evaluated at 24 hpf by gently prodding embryos within the chorion and recording the presence or absence of a twitch response. To minimize user-dependent variability in stimulus force, experiments were repeated by multiple users and the tallies pooled. Hatching rates were assessed at 48 hpf by counting the number of embryos that had successfully hatched versus those remaining within the chorion. Embryos were imaged at the 1-cell stage, 24 hpf, and 48 hpf using an Olympus microscope. Chorion diameter was measured in Fiji by drawing a line across the chorion and converting pixel length to millimeters using a 1 mm microscopic ruler as a calibration standard.

### Single nuclei isolation

Treated embryos and untreated sibling controls were processed together to minimize batch effects. For each condition, 25 dechorionated zebrafish embryos were pooled and flash-frozen. Nuclei were isolated from pooled samples following (Velmeshev et al., 2023) with minor modifications. Briefly, each sample was homogenized in 1 mL RNase-free lysis buffer (0.32 M sucrose, 5 mM CaCl₂, 3 mM MgAc₂, 0.1 mM EDTA, 10 mM Tris-HCl, 1 mM DTT, 0.1% Triton X-100 in nuclease-free water) using a glass Dounce homogenizer (Kontes Glass, Cat. #884900). Homogenates were passed through a 70 µm cell strainer (Celltreat Scientific, Cat. #229483) and transferred to 5 mL open-top Polyallomer ultracentrifuge tubes (Seton Scientific, Cat. #5010). A 1.8 mL sucrose cushion (1.8 M sucrose, 3 mM MgAc₂, 1 mM DTT, 10 mM Tris-HCl in nuclease-free water) was layered beneath each homogenate, and samples were centrifuged at 107,000 × g for 1 h at 4 °C (ThermoFisher Scientific, Sorvall WX Ultra Series, Cat. #46900). Supernatants were carefully aspirated, and the nuclear pellets were incubated in 250 µL nuclease-free 1× PBS for 20 min on ice before resuspension. Nuclear suspensions were filtered through 30 µm sterile strainers (Sysmex Celltric, Cat. #04-004-2326), and nuclei were counted using a hemocytometer. Single-nucleus capture was performed on the 10x Genomics Chromium Controller (Product #1000331) using a target of 10,000 nuclei per sample. Libraries were prepared using the 10x Multiome protocol without modification. Single-nucleus libraries were pooled and sequenced on a NovaSeq 2000 system (Illumina, Cat. #20038897) using P4 flow cells (Cat. #EC1551933-EC14) to an average depth of 20,000 read pairs per nucleus for gene expression libraries and 25,000 read pairs per nucleus for ATAC libraries.

### 10X Multiome Data Analysis

For the initial processing and quality screening to quantify gene expression and chromatin accessibility, a reference genome was constructed using GRCz11 and Ensembl build 105 (Dyer et al., 2025) with CellRanger-arc v2.0.2 (Satpathy et al., 2019; Zheng et al., 2017). Due to suboptimal performance of the ATAC libraries, CellRanger v8.0.1 was used to process the gene expression libraries to filter cells from empty droplets. Further filtering was done by performing k-means clustering of cells within each sample by transcript counts and keeping only cells within the highest-count cluster. Additionally, cells jointly called by CellRAnger-arc were kept. Downstream analysis was performed in Seurat v5 (Hao et al., 2024), using SCTransform (Choudhary & Satija, 2022; Hafemeister & Satija, 2019) for normalization and variance stabilization, reciprocal PCA for integration (Hao et al., 2021), and MAST (Finak et al., 2015) for differential expression tests.

For cell type classification, cell types were setup using the SingleR package (Aran et al., 2019) with the Daniocell atlas (Sur et al., 2023) as a reference and down-sampled to 500 cells per type. For exposure impact on clusters, and to determine the contribution of each cluster to the variance between exposure and control samples, DEGs for each cluster with an adjusted P value <0.05 and a log2 fold change >1.5 were identified and compiled into a matrix of log2 fold changes and used as input for PCA. The contribution of each cluster to the first number of principal components explaining ∼85% of the variance was then calculated. For the cell population proportion test between samples, differential population testing between conditions was carried out using SCProportionTest (Miller et al., 2021).

### HCR probe design and protocol

For florescent in situ Hybridization Chain Reaction (HCR) probes for the top differentially expressed genes were designed using python tool (Kuehn et al., 2022) with details found at Github respository (https://zenodo.org/records/4694867#.ZDcWhuzMKis). Probes for three genes were designed such that they could be multiplexed for visualizing together on the same embryo when doing the HCR protocol. Probes were ordered as oligo pools from IDT and reconstituted with nuclease free water to make a 100uM stock solution.

The HCR procedure was done according to established protocol (Choi et al., 2018) provided by Molecular Instruments with minor modifications. Briefly, zebrafish embryos were dechorionated and fixed overnight in 4% paraformaldehyde at 4 °C, then dehydrated through a methanol series and stored at −20 °C until use. All untreated and treated samples were processed and imaged together to minimize batch effects. Prior to hybridization, samples were rehydrated through a methanol series into 1× PBST. Embryos were then treated with 2 ng/L Proteinase K for 5 min, followed by PBST washes. Samples were transferred to mesh baskets and incubated in 200 µL pre-hybridization solution for 30 min at 37 °C. Hybridization solution containing probes was then added (1 µL probe stock per 100 µL hybridization solution). Up to three probes were applied simultaneously, and samples were incubated overnight at 37 °C. The following day, hybridization solution was removed and samples were washed with wash buffer and 1× SSCT. Amplification buffer was added, and samples were incubated at room temperature for 30 min. Hairpins corresponding to the selected probes were prepared by adding 2 µL each of h1 and h2 hairpin stock per 100 µL amplification buffer, allowing up to three fluorophore-conjugated hairpins per sample. Hairpin mixtures were denatured at 95 °C for 90 s and allowed to cool in the dark at room temperature before being added to samples. Samples were then incubated overnight at room temperature in the dark. The next day, hairpins were removed and samples were washed with 1× SSCT. Nuclei were counterstained with DAPI using NucBlue (Catalog number R37606), 1 drop per 1,000 µL 1× SSCT, for 20 mins at room temperature, followed by incubation at 4 °C for 2 h. Samples were imaged on a Zeiss confocal LSM 800 microscope using 488, 546, and 647 nm lasers at 20× magnification, with identical laser power settings across all treatment conditions. Image analysis was performed using Imaris software and its deep-learning pipeline to quantify either bright spots corresponding to nascent transcription or gross signal representing processed single transcripts across defined regions of interest.

### Behavioral analysis

Zebrafish embryos were exposed to fresh PFOA-containing media daily from the 1-cell stage through 5 dpf while maintained in Petri dishes. At 5 dpf, larvae were randomly selected and transferred to a 24-well plate, with one larva per well. Larval behavior was assessed using a DanioVision system. Movement was recorded for 1 min under baseline conditions without stimulus, followed by 1 min with a light stimulus and 1 min with a mechanical stimulus (plate movement). Total distance moved was recorded for each condition. Each well was divided into a central zone (zone 1) and a peripheral zone (zone 2), and distance traveled within each zone was quantified. Behavioral tracking and data extraction were performed using EthoVision software. Statistical outputs from EthoVision were used to generate plots in RStudio using the ggplot package.

