## Supplementary material for "Early exposure to PFAS disrupts neuro-muscular development in zebrafish embryos": Supp Figure

**List of supplementary figures**

Supp figure 1 – tail twitch poke video
Supp figure 2 – chorion size differences at even half an hour after treatment
Supp figure 3 – bar graph of most impacted tissue clusters (PC.contib – geX folder)
Supp figure 4-5 – heatmaps for muscle and neuronal clusters (heatmap pdfs folder)
Supp figure 6 – Changes in cell counts for tissue cluster (cluster differences folder)
Supp figure 7 – sox6 binding sites with pcdh7b genomic loci
Supp figure 8 – foxp4 forebrain expression
Supp figure 9 – TF motifs present in pcdh7b altered chromatin regions
Supp figure 10 – TF motifs present in *sox6* altered chromatin regions
Supp figure 11 – TF motifs present in *foxp4* altered chromatin regions
Supp figure 12 – behavior heatmap and graphs for 5 dpf

Supp Fig 1

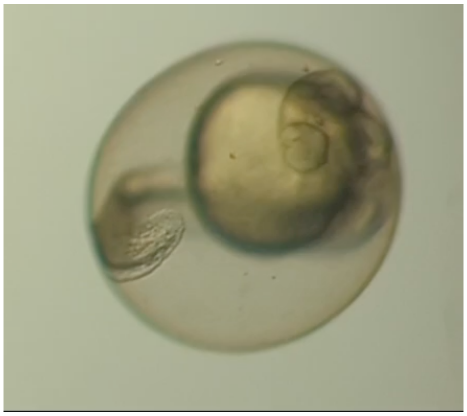

Supp figure 2

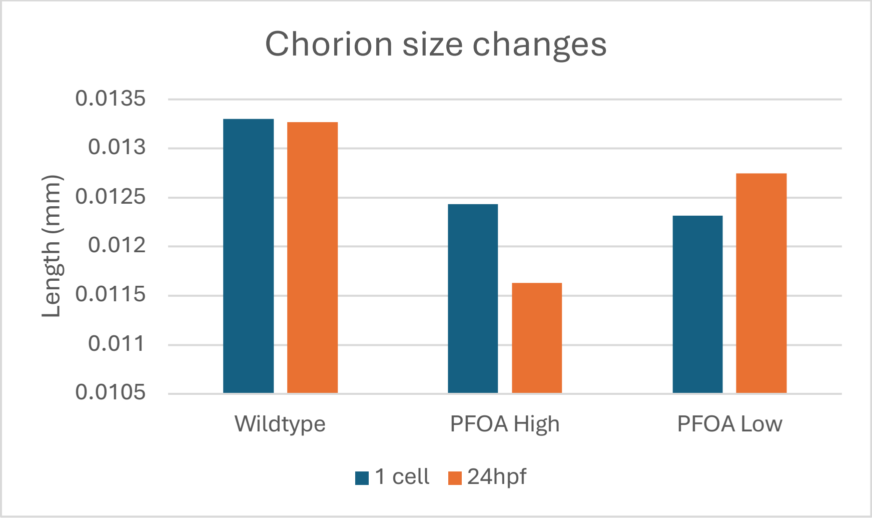

### Supp figure 3

#### Supp fig 3

##### Comparing PFOA low exposure to untreated

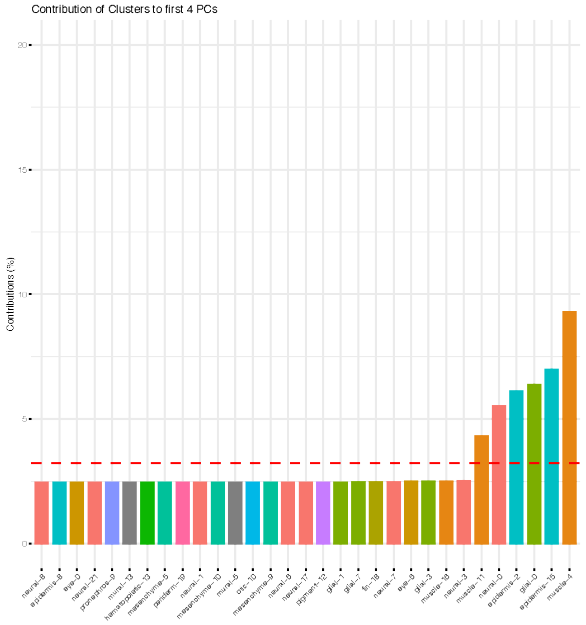


##### Comparing PFOA high exposure to untreated

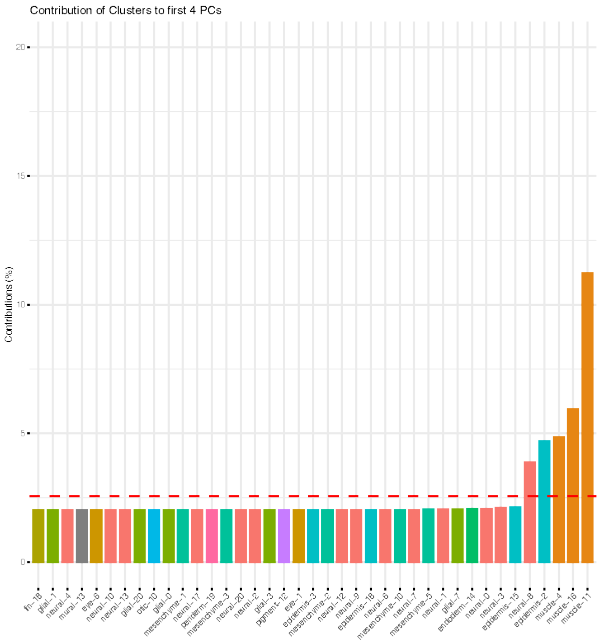


##### Comparing PFOA parental exposure to untreated

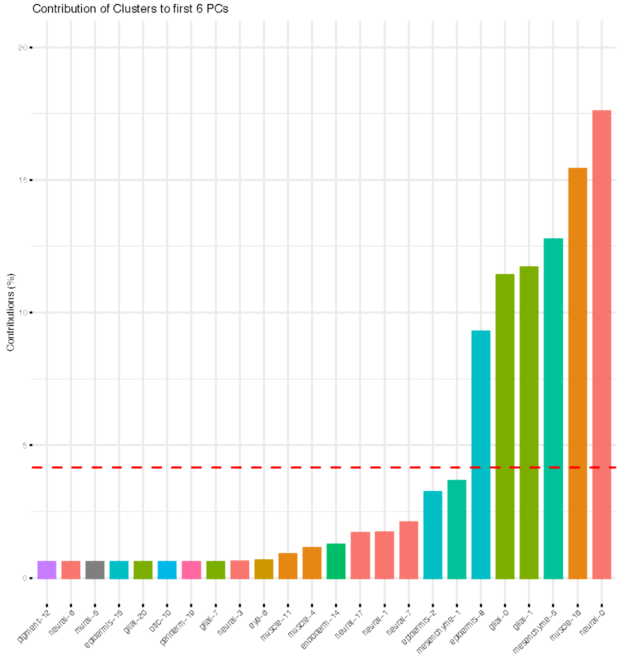

### Supp figure 4

#### Comparing low PFOA exposure to untreated (none)

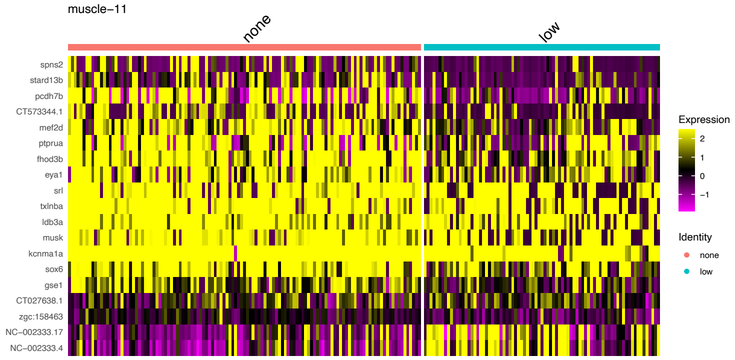


#### Comparing high PFOA exposure to untreated (none)

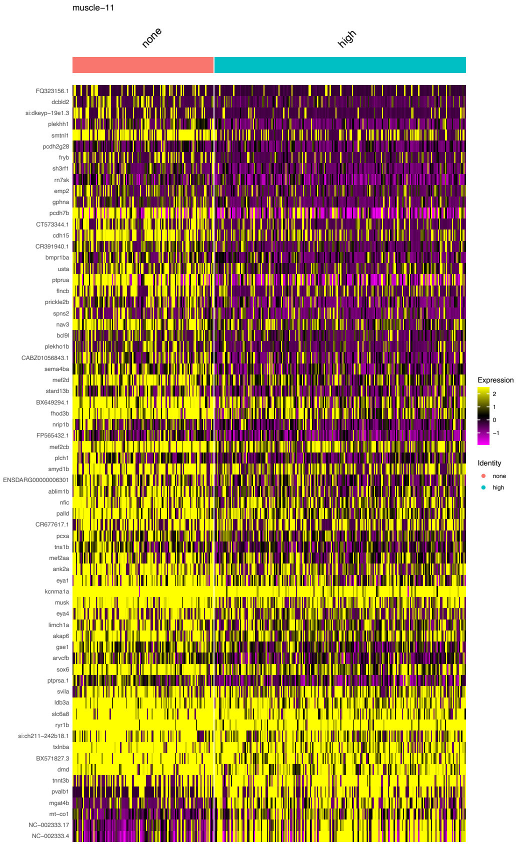


#### Comparing parental PFOA exposure to untreated (none)

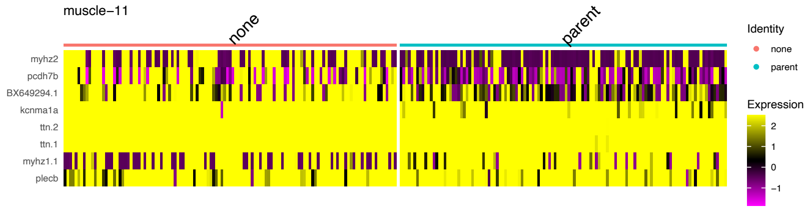

### Supp fig 5

#### Comparing low PFOA exposure to untreated (none)

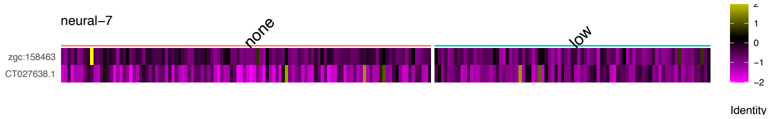


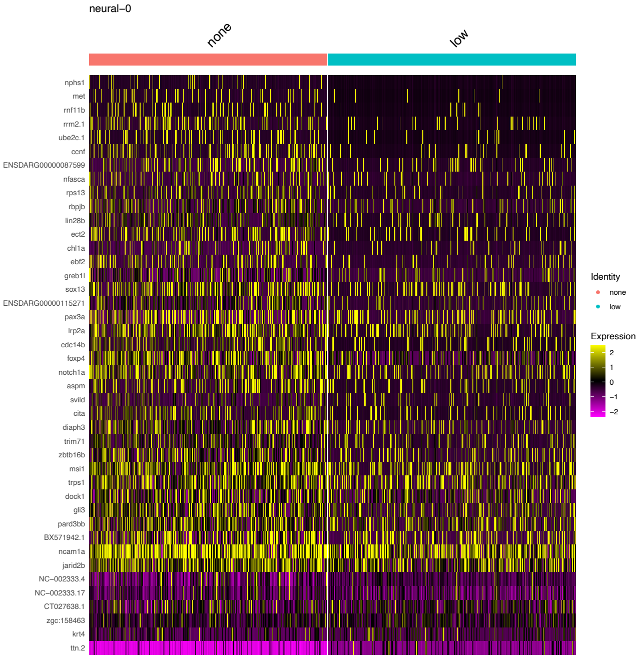


#### Comparing high PFOA exposure to untreated (none)

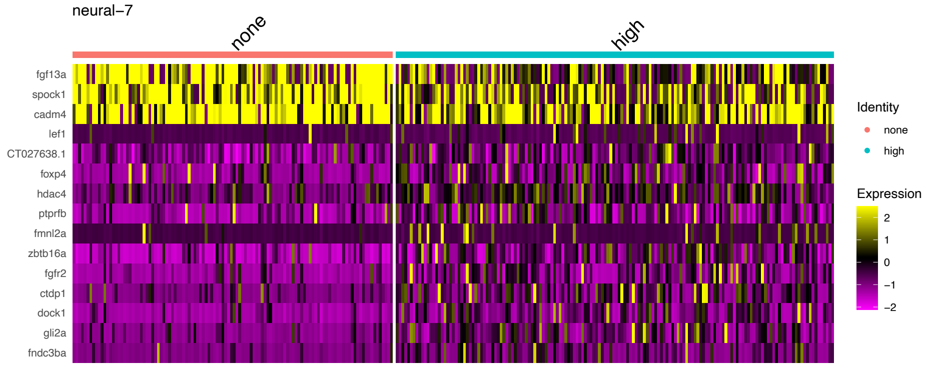


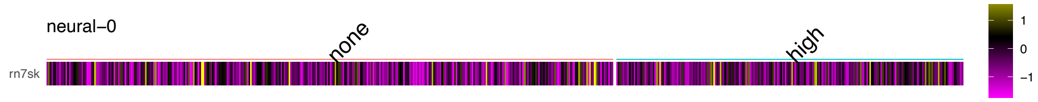


#### Comparing parental PFOA exposure to untreated (none)

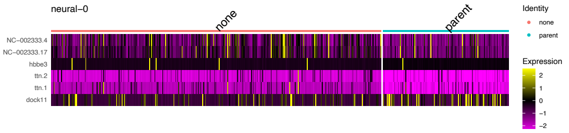


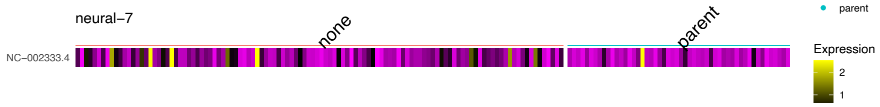

Supp figure 6

#### Comparing low PFOA exposure to untreated (none)

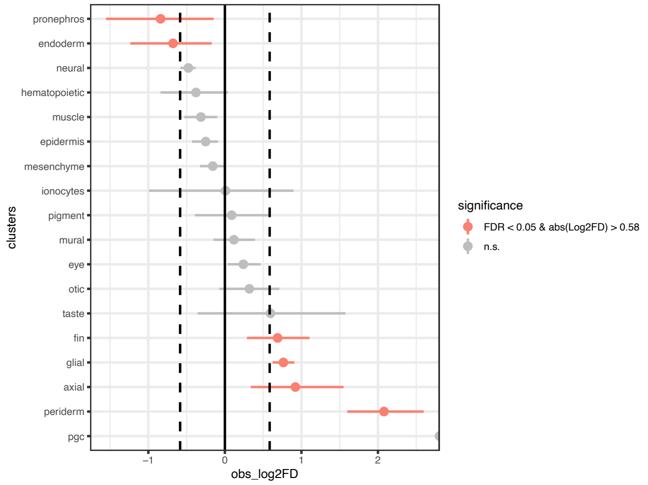


#### Comparing high PFOA exposure to untreated (none)

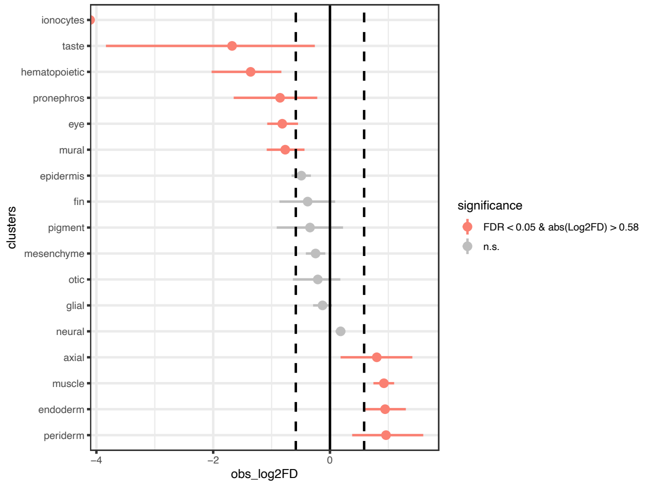


#### Comparing parental PFOA exposure to untreated (none)

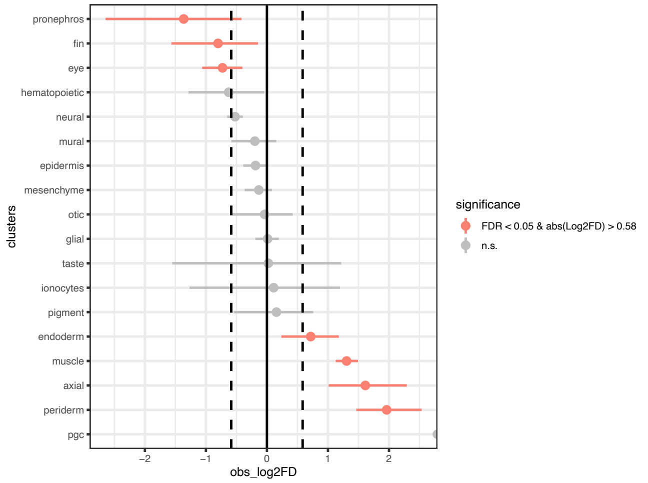

Supp figure 7

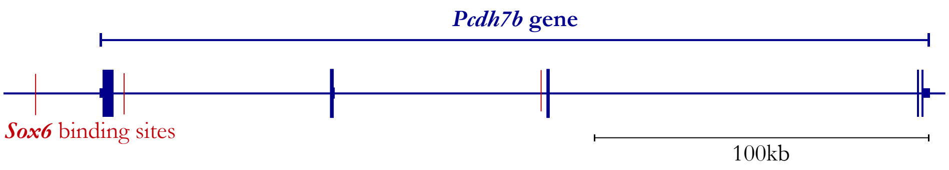


Supp figure 8

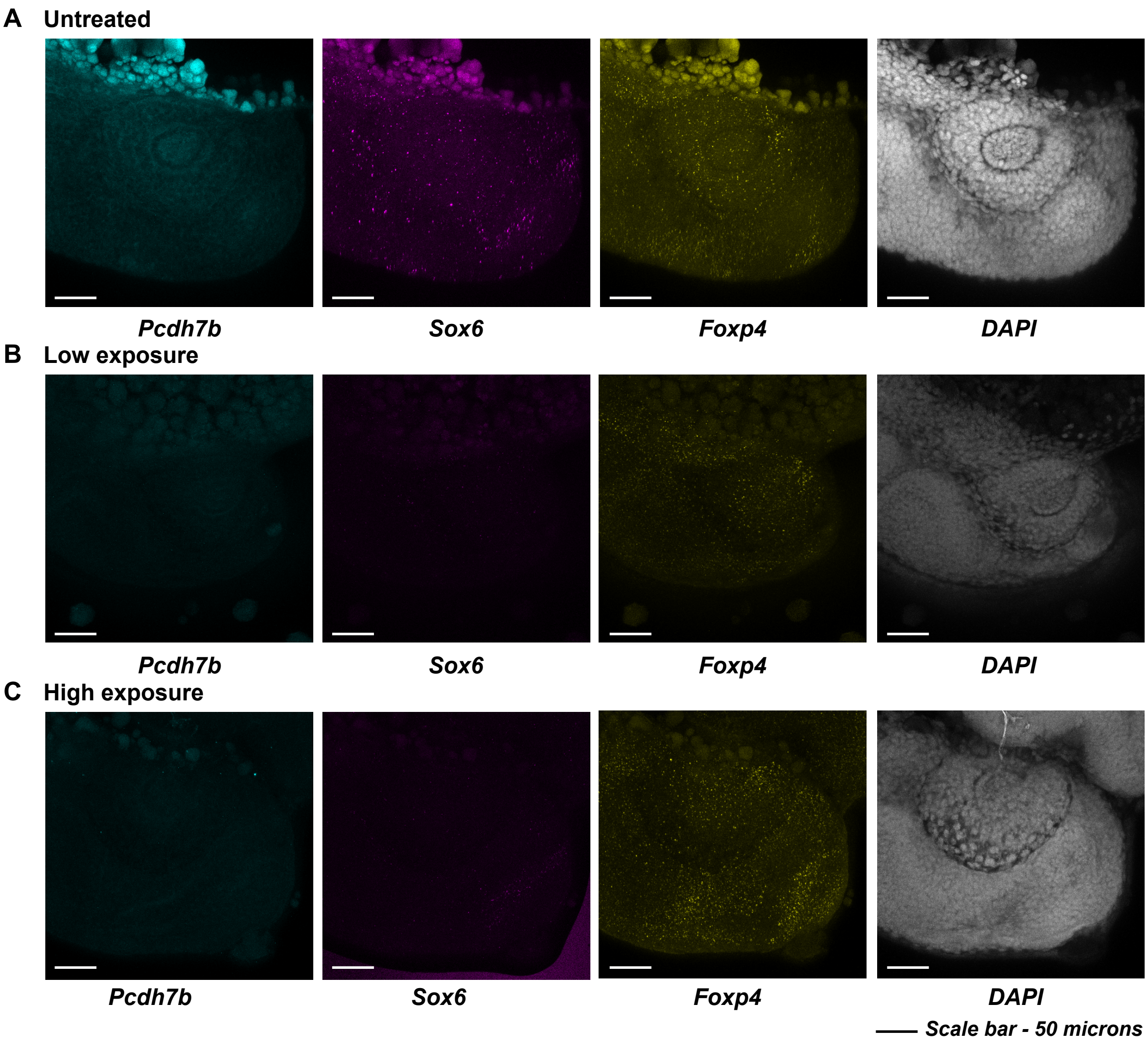

Supp figure 9 – TF motifs present in pcdh7b altered chromatin regions

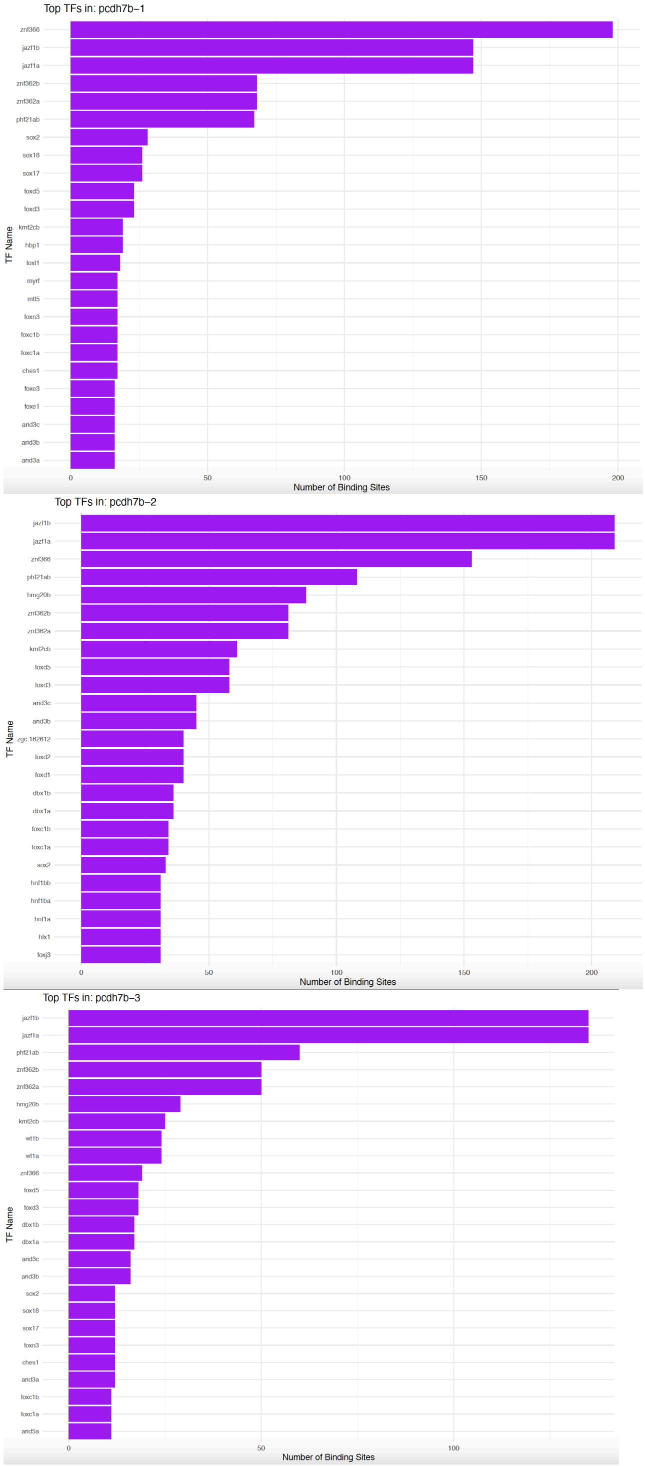

Supp figure 10 – TF motifs present in *sox6* altered chromatin regions

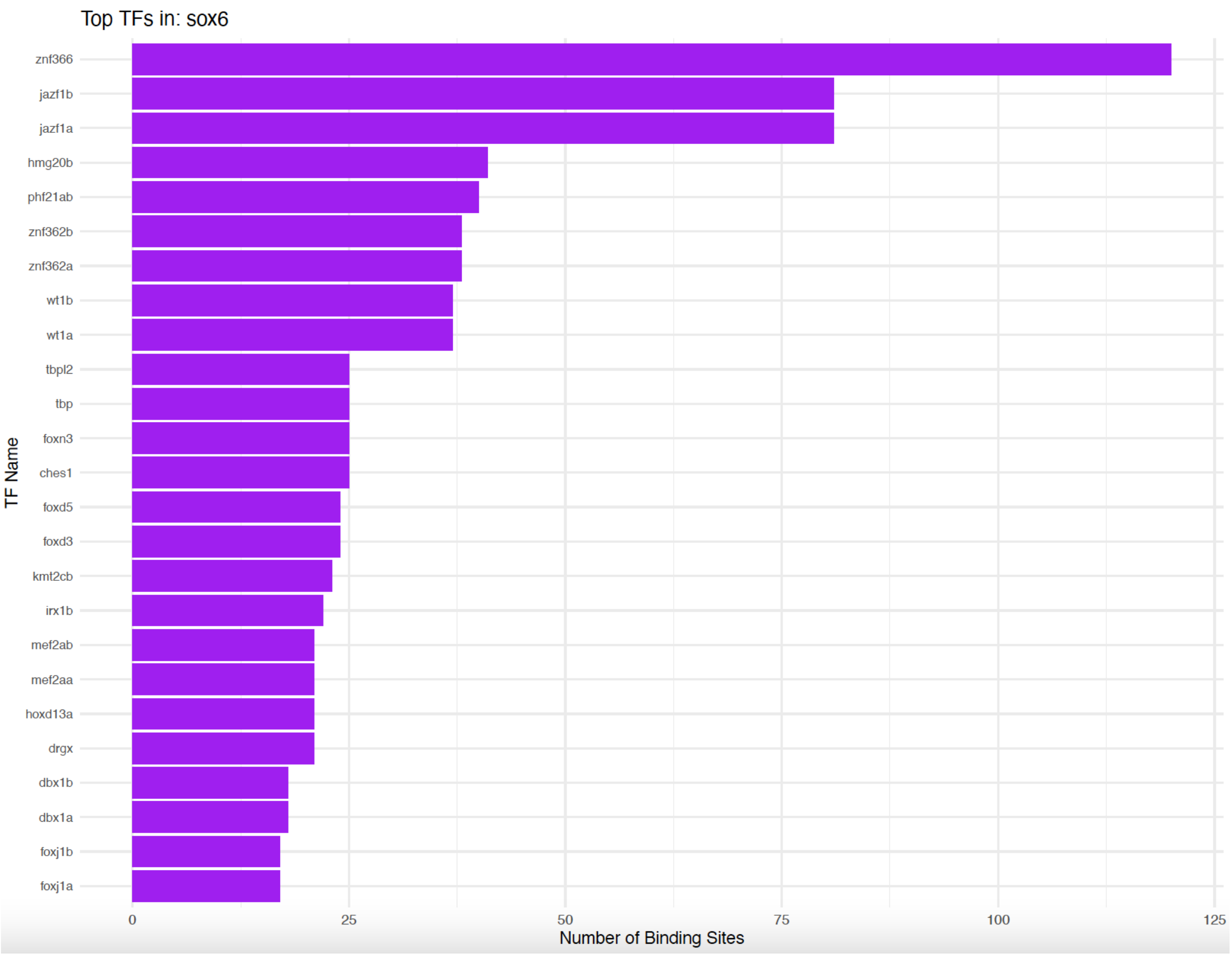

Supp figure 11 – TF motifs present in *foxp4* altered chromatin regions

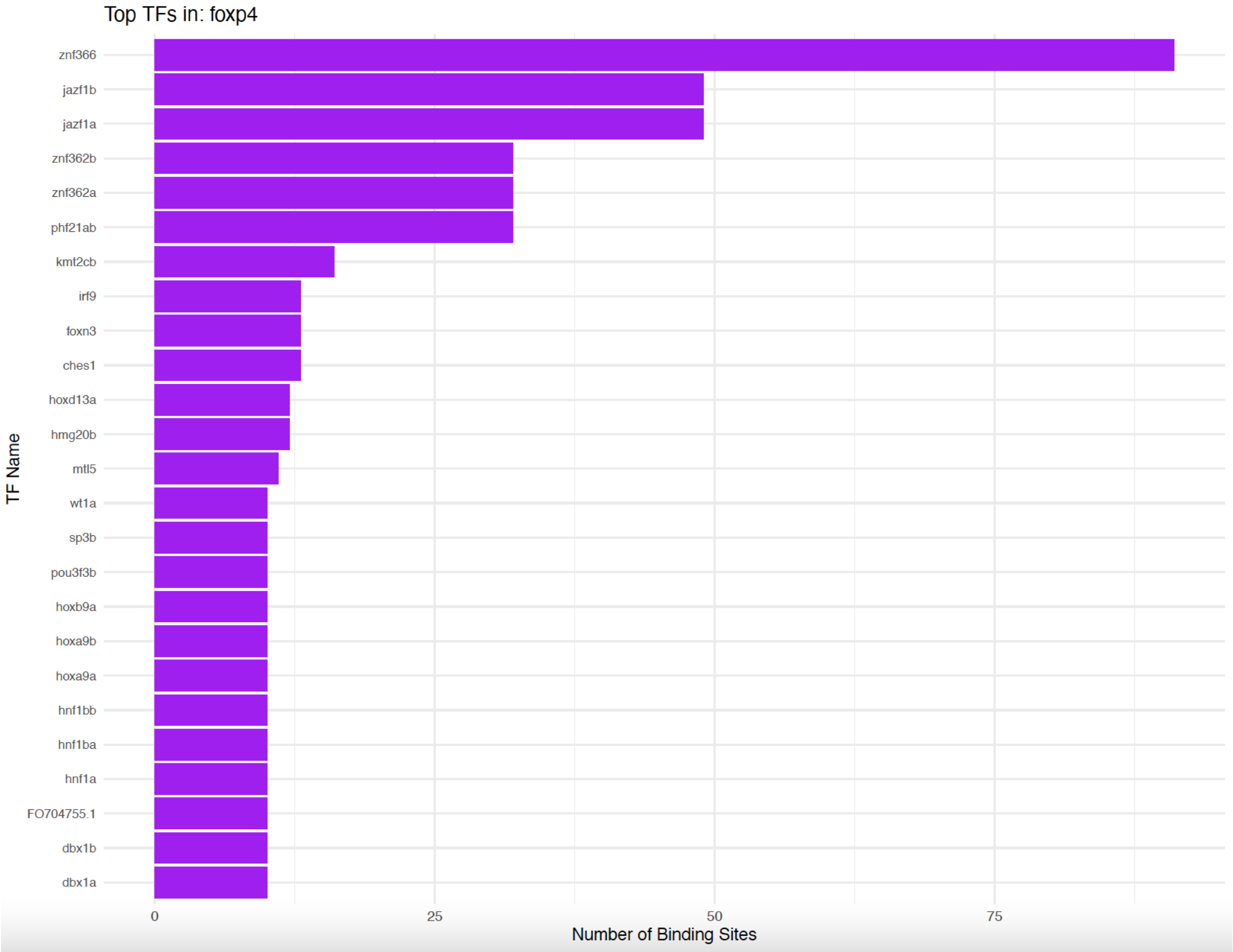

Supp figure 12

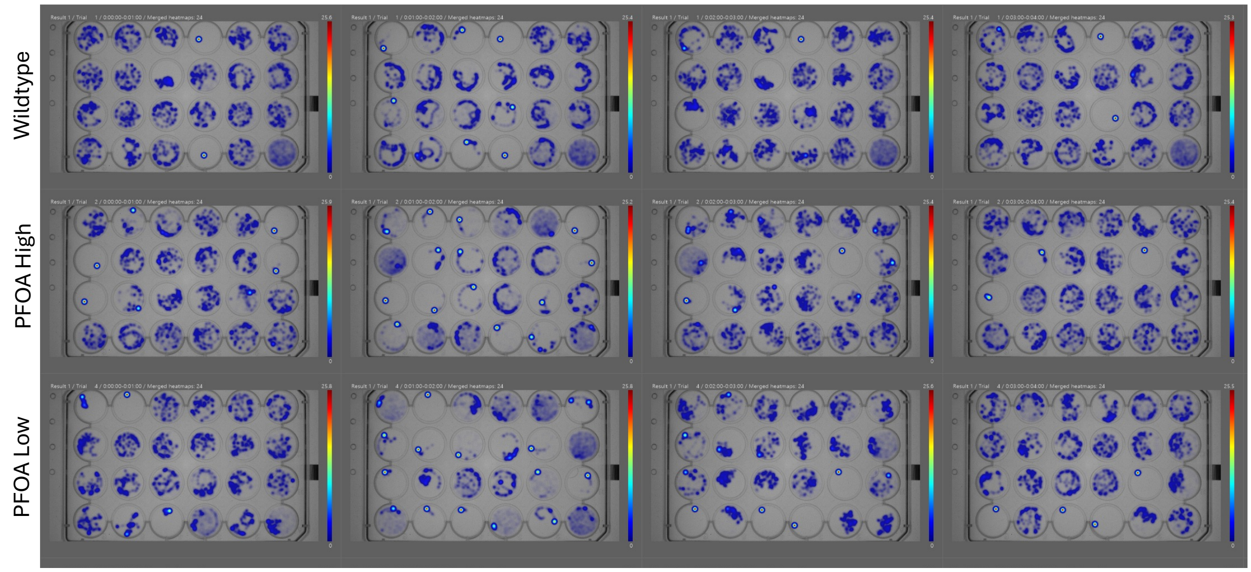


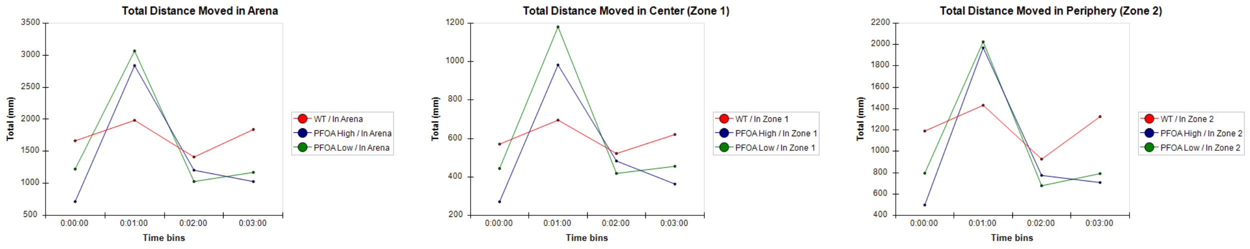


Zone 1 – center
Zone 2 – periphery
Arena – whole well

Time 0-1 mins – no stimulus
Time 1-2 mins – light on
Time 2-3 mins – light off
Time 3-4 mins – trigger intensity 4
